# Bioinformatics Approaches to Determine the Effect of SARS-CoV-2 Infection on Patients with Intrahepatic Cholangiocarcinoma

**DOI:** 10.1101/2023.01.04.522709

**Authors:** Xinyi Zhou, Tengda Huang, Hongyuan Pan, Jiang Lan, Tian Wu, Ao Du, Yujia Song, Yue Lv, Kefei Yuan

## Abstract

Severe acute respiratory syndrome coronavirus 2 (SARS-CoV-2), the causal agent of coronavirus disease 2019 (COVID-19), has infected millions of individuals throughout the world, which poses a serious threat to human health. COVID-19 is a systemic disease that affects tissues and organs, including the lung and liver. Hepatocellular carcinoma (HCC) and intrahepatic cholangiocarcinoma (ICC) are the most common liver cancer, and cancer patients are particularly at high risk of SARS-CoV-2 infection. The relationship between HCC and COVID-19 has been reported in previous studies, but ICC has been rare. With the methods of systems biology and bioinformatics, this study explored the link between COVID-19 and ICC. Transcriptional profiling of COVID-19 and ICC were obtained from the GEO database. A total of 70 common differentially expressed gene (DEGs) of both diseases were identified to investigate shared pathways. Then top-ranked 10 key DEGs (SCD, ACSL5, ACAT2, HSD17B4, ALDOA, ACSS1, ACADSB, CYP51A1, PSAT1, and HKDC1) were identified as hub genes by protein-protein interaction (PPI) network analysis. In addition, transcriptional regulatory networks regulating hub genes were revealed by hub Gene-transcription factor (TF) interaction analysis and hub gene-microRNA (miRNAs) interaction analysis. This study is expected to provide new references for future research and treatment of COVID-19 and ICC.

## 1 Introduction

Coronavirus disease 2019 (COVID-19) was recognized in December 2019(1), which was a recently discovered respiratory condition brought on by severe acute respiratory syndrome coronavirus 2 (SARS-CoV-2). Most COVID-19 patients have mild to moderate symptoms, but 5% of them also suffer acute respiratory distress syndrome (ARDS), multiple organ failure, or septic shock, and around 15% of patients develop severe pneumonia(2). New SARS-CoV-2 variants continue to emerge (such as Alpha, Beta, Delta and Omicron) and are associated with high case rates and substantial global mortality. As of December 2022, there have been 650,332,899 cases of COVID-19, resulting in 6,649,874 deaths, according to the World Health Organization (WHO). The earlier research reveal SARS-CoV-2 infection mostly happens when the surface spike protein of the virus binds to the angiotensin-converting enzyme 2 (ACE2) receptor on human cells. The spike protein is the protein for SARS-CoV-2 to recognize host cells and is also the main target of the human immune system(3). Vaccines against SARS-CoV-2 are being developed and tested at an unprecedented rate. So far, various vaccines against COVID-19 have been produced, including those created by Pfizer-BioNTech(4), Sinovac Biotech(5), and AstraZeneca-University of Oxford(6). However, effective medications to treat and prevent COVID-19 are in scarcity.

Although the virus directly infects the lungs, its effect on the liver cannot be ignored. The data from The Fifth Medical Center of PLS General Hospital, Beijing, China indicate that 2-11% of patients with COVID-19 had liver comorbidities and 14-53% cases reported abnormal levels of alanine aminotransferase (ALT) and aspartate aminotransferase (AST) during disease progression. And patients with severe COVID-19 seem to have higher rates of liver dysfunction(7). Liver cancer is the sixth most common and the third deadliest malignancy in the world(8). The relationship between COVID-19 and hepatocellular carcinoma (HCC), the most common form of liver cancer, has been reported recently(9). As the second most common liver cancer, ICC is highly similar with HCC, which are both associated with cirrhosis, infection of Hepatitis B Virus (HBV) and Hepatitis C Virus (HCV), and metabolic syndrome(10). First-line (gemcitabine and cisplatin), second-line (FOLFOX), and adjuvant (capecitabine) systemic chemotherapy are currently the accepted standard of treatment of ICC(11). In fact, cancer patients treated by chemotherapy or immunotherapy are more susceptible to COVID-19 infection(12). In addition, ICC patients had more severe liver dysfunction. Therefore, to better overcome COVID-19 and ICC in the future, it is imperative to explore and clarify the internal molecular mechanism between these two diseases.

Two datasets were utilized in this work to explore the correlations between ICC and COVID-19. GSE152418 and GSE119336 for COVID-19 and ICC, respectively, were acquired from the Gene Expression Omnibus (GEO) database. Initially, differentially expressed genes (DEGs) were identified in the data set, and then 70 shared DEGs genes were found for both diseases. Pathway analysis was done using these mutual DEGs to better study the mechanisms based on gene expression. To gather hub genes, the network of protein-protein interactions (PPIs) is also created using the 70 recognized DEGs. The hub genes are the major experimental genes for the following investigation. The extraction of hub genes from shared DEGs is a critical step in the building of gene regulation networks. Next, the hub genes were used to delineate the gene-regulatory network and complete the gene-disease association network. A flowchart of the overall work is presented in Figure 1. Our analysis reveals a pathway for co-treatment of the two diseases.

**Figure 1.**
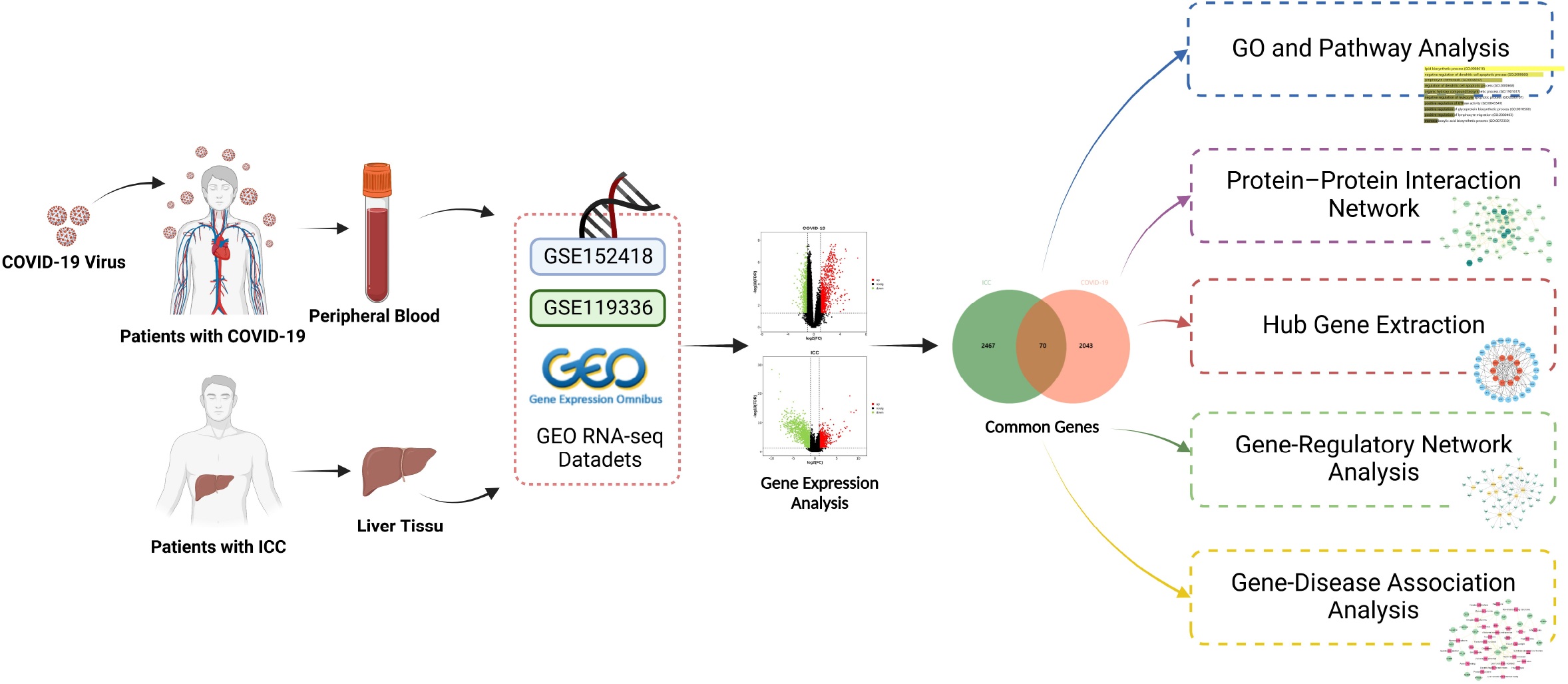
Schematic illustration of the overall general workflow of this study.

## 2 Materials and Methods

### 2.1 Collection of Gene Expression Data

RNA-seq datasets from the National Center for Biotechnology Information (NCBI)(12), GEO (http://www.ncbi.nlm.nih.gov/geo) database were obtained to investigate common biological interrelationships between COVID-19 and ICC. The GEO accession ID of the COVID-19 dataset is GSE152418, which is transcriptional profiling of peripheral blood from 17 healthy individuals and 17 COVID-19 infected individuals through high-throughput sequencing by Illumina NovaSeq 6000 (Homo sapiens) platform for extracting RNA sequence(13). The ICC dataset (GEO accession ID: GSE119336) contains 15 pairs of human ICC tumors and matched non-tumor liver tissues that were sequenced using an Illumina HiSeq 2000 (Homo sapiens) high-throughput sequencing machine donated by Zhou et al(14).

### 2.2 Identification of DEGs and Shared DEGs between ICC and COVID-19

If a significant difference in statistics between the various test conditions at the transcriptional level, genes are classified as DEGs(15). LIMMA package(16) was adopted to identify the DEGs of GSE119336 and GSE152418 from the expression values, and was corrected by Benjamini-Hochberg to reduce the false discovery rate (FDR). Cutoff criteria (FDR < 0.05 and |log_2_FC| > 1) was used to perceive significant DEGs from both datasets. The shared DEGs of GSE119336 and GSE152418 were acquired using the jvenn(17) (http://jvenn.toulouse.inra.fr/app/example.html), an online VENN graph mapping platform, to plot VENN analysis.

### 2.3 Gene Ontology (GO) and Pathway Enrichment Analysis

An important analytical endeavor aiming at biologically classified genes is gene enrichment analysis(18). In order to understand the function of common DEGs, we performed GO and pathway enrichment analyses connected with the mutual DEGs using Enrichr(19) (https://maayanlab.cloud/Enrichr/), a wide range of online gene set enrichment tool. The three types of GO database in the GO database are biological processes, molecular functions, and cellular components. In pathway enrichment analysis, four databases were regarded, including Kyoto Encyclopedia of Genes and Genomes (KEGG) 2021 human pathway, Wikipathway 2021, Reactome 2022, and the Bioplanet 2019. And *P* < 0.05 was used as a criterion to screen for reliable results.

### 2.4 PPI Network Analysis

The PPI was identified using the STRING(20) (version 11.5) database (https://cn.string-db.org/), an online protein-protein association networks platform, and was then visualized and drawn as a network using Cytoscape(21) (version 3.9.1), an open source software platform for visualizing complex networks. Using a composite score larger than 0.15 and proteins encoded by the common DEGs between COVID-19 and ICC, the PPI network was constructed. Hub genes are genes that exhibit strong connections in potential modules(22). The PPI network modules were then ranked, examined, and hub genes were predicted using cytoHubba, a Cytoscape plug-in.

### 2.5 Gene-Regulatory Network Analysis

To discover the transcriptional factors (TFs) and microRNAs (miRNAs) that regulate the hub genes post-transcriptionally, hub gene-TF interplay networks and hub gene-miRNA interaction networks have been dug by means of NetworkAnalyst(23) (version 3.0), a comprehensive visual analysis platform for gene expression profiling. The hub gene-TF interaction networks were built according to the JASPAR(24) database. Hub gene-miRNA interaction networks were constructed via the TarBase(25) (version 8.0) databases.

### 2.6 Gene-Disease Association Analysis

In order to study the human genetic illnesses of shared genes between COVID-19 and ICC, DisGeNET(26) (https://www.disgenet.org/), one of the largest publicly accessible databases of genes and variations linked to human illnesses may be found on a discovery platform, was used in our analysis. Currently, DisGeNET has information on about 24,000 illnesses and features, 17,000 genes, and 117,000 genetic variations(26). Similarly, NetworkAnalyst and Cytoscape were used to dig gene-disease relationships in order to find diseases associated with common DEGs.

## 3 Result

### 3.1 Recognition of DEGs and biological relationships between ICC and COVID-19

Our work examined the human RNA-seq dataset from the NCBI to discover DEGs of COVID-19 and ICC in order to evaluate the interactions and implications of ICC with COVID-19. In the ICC dataset, our analysis found 2,537 DEGs, of which 1,095 were up-regulated and 1,442 were down-regulated (Figure 2A, Supplementary Table 1). Meanwhile, 2,113 genes showed differential expression in the COVID-19 dataset, including 1,267 up-regulated and 891 down-regulated genes (Figure 2B, Supplementary Table 2). Table 1 is a list of the condensed data of DEGs. With the use of the cross-comparative analysis, we were able to find 70 DEGs that were shared by the ICC and COVID-19 datasets (Figure 2C, Supplementary Table 3). This common gene set is used to carry out further analyses. The outcomes of differential expression analysis revealed that COVID-19 and ICC had certain molecular similarities and interacted in certain ways.

**Figure 2.**
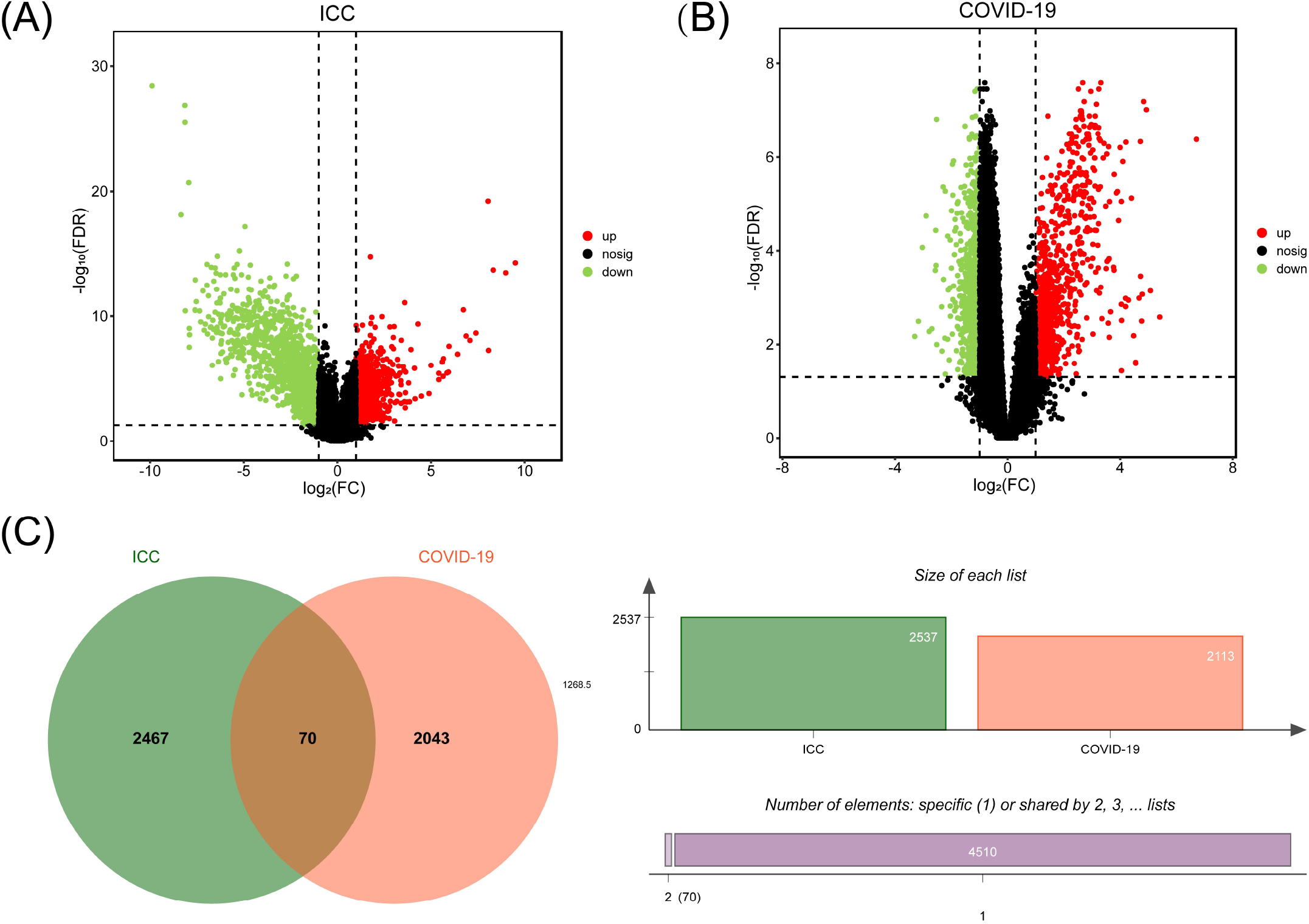
Volcano plots of (A) COVID-19 and (B) ICC, with genes with |log2FC| > 1 and FDR < 0.05. (C) The Venn diagram depicts the shared DEGs among COVID-19 and ICC.

**Table 1.** Overview of datasets with their geo-features and their quantitative measurements in this analysis.

| Disease name | GEO accession | GEO platform | Total DEGs count | Up regulated DEGs count | Down regulated DEGs count |
| --- | --- | --- | --- | --- | --- |
| ICC | GSE119336 | GPL11154 | 2,537 | 1,095 | 1,442 |
| COVID-19 | GSE152418 | GPL24676 | 2,113 | 1,267 | 846 |

### 3.2 DEGs Functional Enrichment Analysis Identified Important Terms and Pathways

Our study used GO and pathway enrichment analysis to learn more about the roles and signaling pathways these typical DEGs play. Gene functional similarity is frequently assessed using the GO enrichment analysis(27). A modeling technique called pathway analysis is used to show how crucial molecular or biological processes interact and illustrate the reciprocal impacts of various diseases(28). In order to uncover highly enriched functional GO keywords and pathways, we ran a functional-enrichment test on common DEGs using the Enrichr program.

70 common DEGs were enriched in 334 terms, including 253 biological processes, 65 molecular functions, and 16 cellular components (Supplementary Table 4). Then we summarized the top 10 terms according to *P*-value in each category in Table 2 and visualized in Figure 3(A-C). It can be found that many of these terms are related to metabolism and immunity, such as lipid biosynthetic process (GO: 0008610) and negative regulation of dendritic cell apoptotic process (GO:2000669), which have a strong association with COVID-19 and ICC.

**Figure 3.**
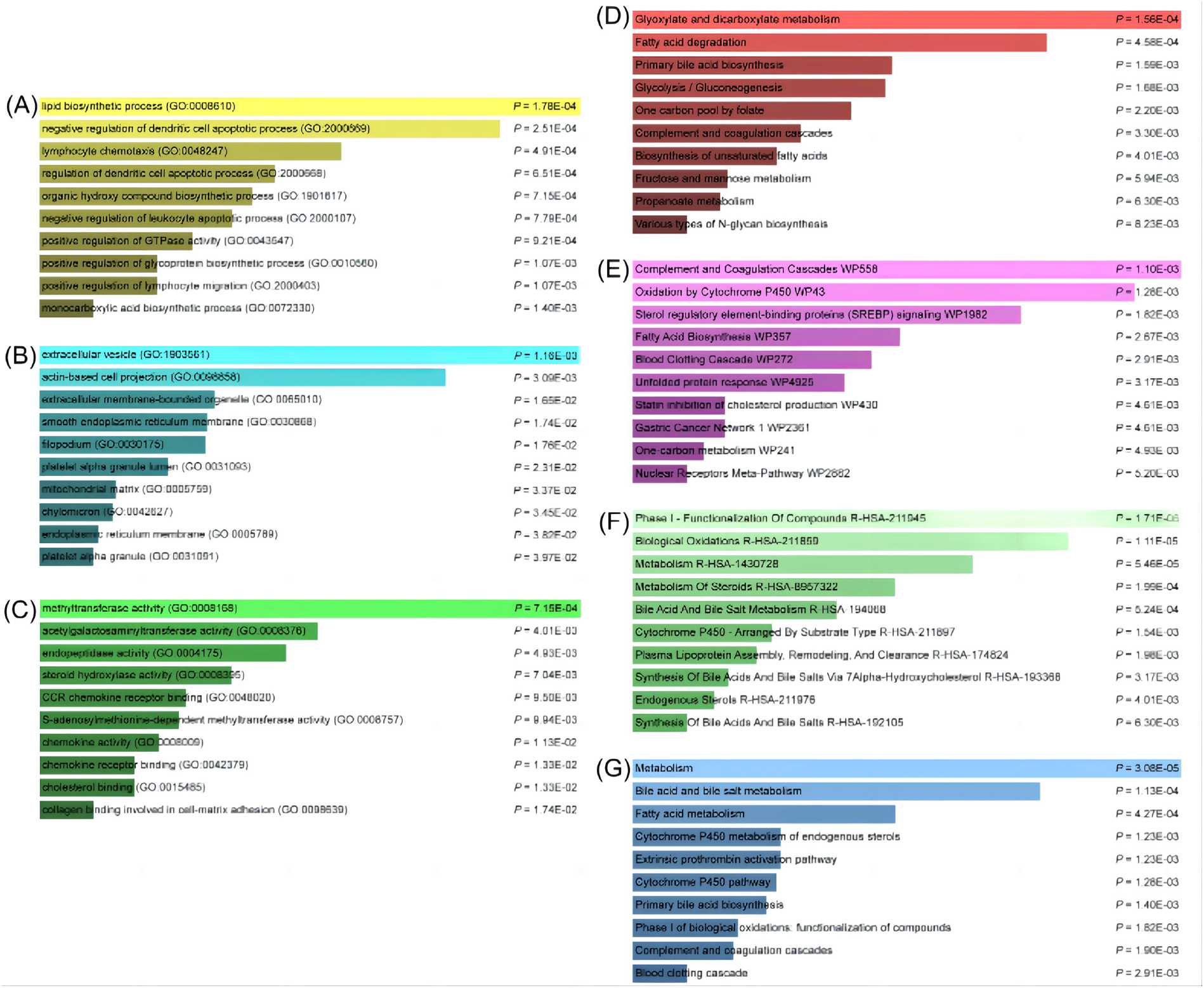
Analysis of common DEGs between COVID-19 and ICC using ontology and pathway enrichment. Ontological analysis: (A) biological processes, (B) molecular function, and (C) cellular components. Pathway enrichment analysis: (D) KEGG 2021 human pathway, (E) Wikipathway 2021, (F) Reactome 2022 pathway, and the (G) Bioplanet 2019 pathway.

**Table 2.**
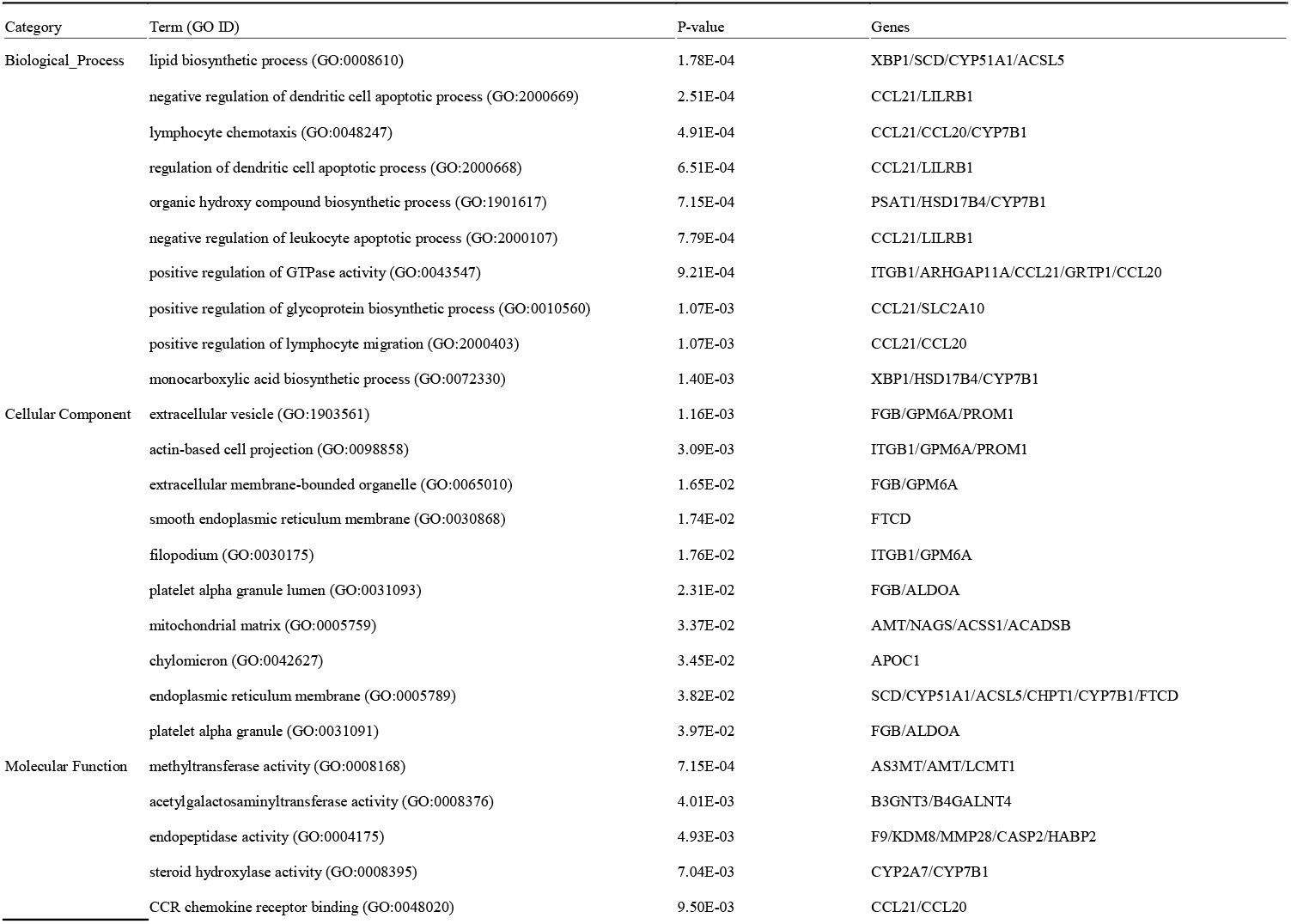
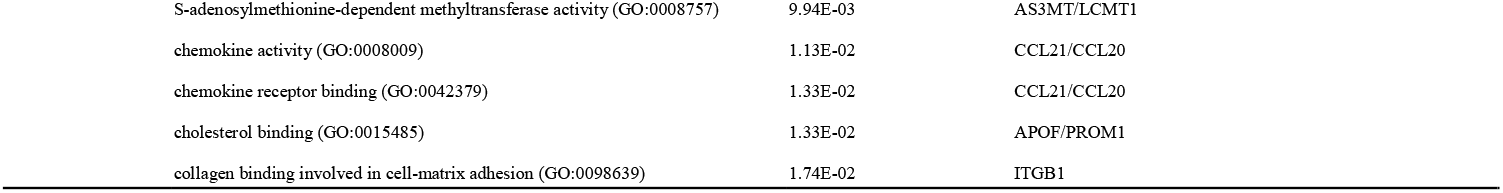
Ontological analysis of common DEGs between COVID-19 and ICC.

We found 22 reliable pathways in KEGG, 32 reliable pathways in Wikipathway, 62 reliable pathways in Reatcome, and 77 reliable pathways in Bioplanet (Supplementary Table 5). The top 10 reliable pathways found in each database are listed in Table 3, and the bar graphs of pathway enrichment analysis are shown in Figure 3(D-G). In these pathways, more about the metabolic pathways were discovered, such as glyoxylate and dicarboxylate metabolism in KEGG, fatty acid biosynthesis in Wikipathway, metabolism of steroids in Reactome, and bile acid and bile salt metabolism in BioPlanet, which indicated that COVID-19 and ICC have common effects on these pathways.

**Table 3.**
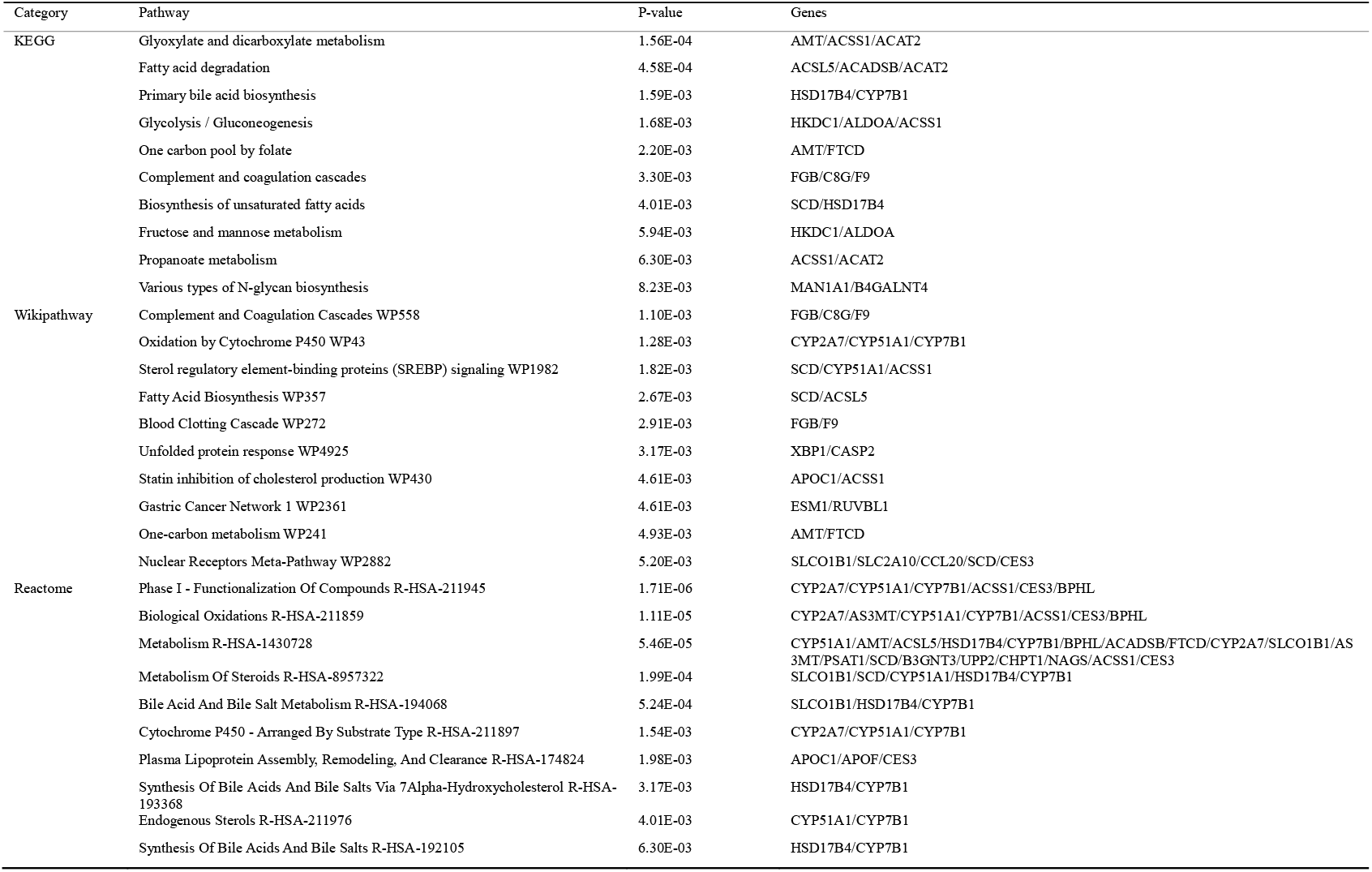
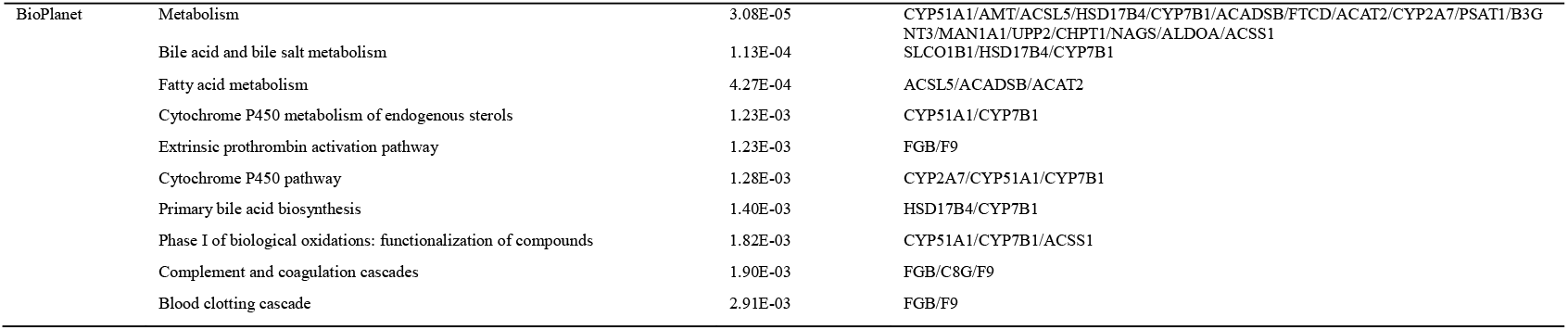
Pathway enrichment analysis of common DEGs between COVID-19 and ICC.

### 3.3 PPI Networks Analysis and Identification of Hub Genes

To better understand biological signals, response mechanisms of energy substance metabolism, and functional links between proteins in disease states, this study obtained the PPI network via STRING. Subsequently, PPI was visualized in Cytoscape to forecast interaction between common protein-coding DEGs. The PPI network of common DEGs consists of 65 nodes and 177 edges (Figure 4, Supplementary Table 6). According to PPI network analysis integrating Cytohubba plugin in Cytoscape, we ranked the most interconnected nodes top 10 DEGs (14.28%) as hub genes. The hub genes are as follows: *SCD, ACSL5, ACAT2, HSD17B4, ALDOA, ACSS1, ACADSB, CYP51A1, PSAT1*, and *HKDC1*. With the aid of the Cytohubba plugin, we also built a network of submodules to better comprehend their closeness and close connection, including 35 nodes and 114 edges (Figure 5). In the following analysis, we will focus on these 10 hub genes. These hub genes show potential biomarkers that can provide new therapeutic strategies for COVID-19 and ICC.

**Figure 4.**
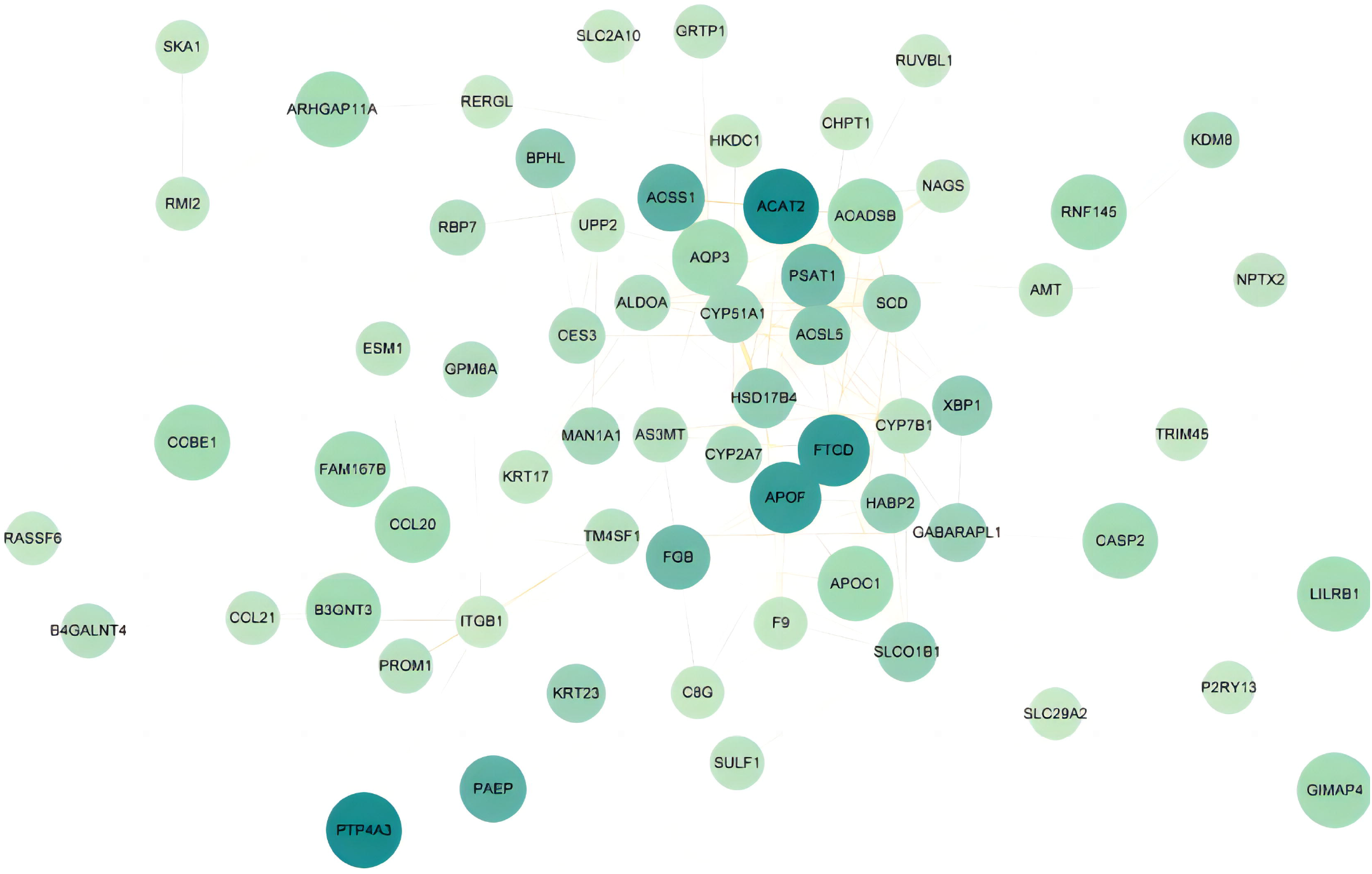
PPI network of common DEGs between COVID-19 and ICC. The circular nodes in the figure stand in for DEGs, while the edges indicate node interactions. The PPI network consists of 177 edges and 65 nodes. String was used to create the PPI network, and Cytoscape was used to display it.

**Figure 5.**
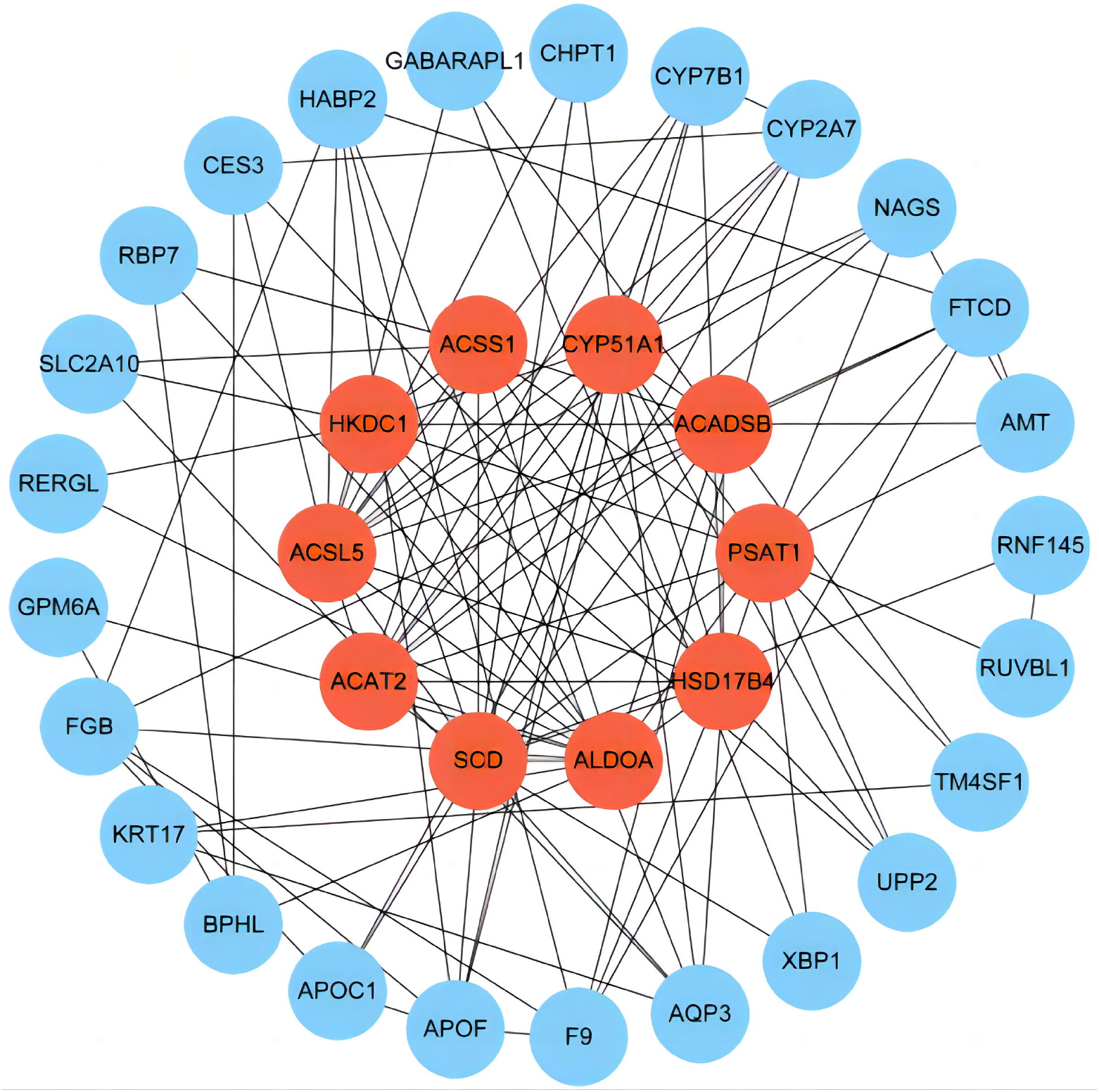
Determination of hub genes from the PPI network by using the Cytohubba plugin in Cytosacpe. Hub genes were obtained using the Cytohubba plugin. Here, the red nodes indicate the highlighted top 10 hub genes and their interactions with other molecules. The network consists of 35 nodes and 144 edges.

### 3.4 Construction of Regulatory Networks at Transcriptional Level

To better understand the regulatory hub genes and detect the key alterations at the transcriptional level, we used network analysis to search for TFs and miRNAs of regulatory hub genes. The TFs-hub genes interactions are shown in Figure 6, and the information of interaction is presented in Supplementary Table 7. 44 TFs were found in the network, and *ALDOA, CYP51A1, ACSL5, SCD*, and *CREB1* were more highly expressed among hub genes as genes have a higher degree in the network of TF-hub gene interactions. Figure 7 and Supplementary Table 8 depict the relationships of miRNA-hub genes. By the similar method, multiple discovered hub genes were projected to be regulated by 112 miRNAs, such as *SCD, ALDOA, PSAT1, CYP51A1*. In-depth study of these genes has common implications for the treatment of COVID-19 and ICC.

**Figure 6.**
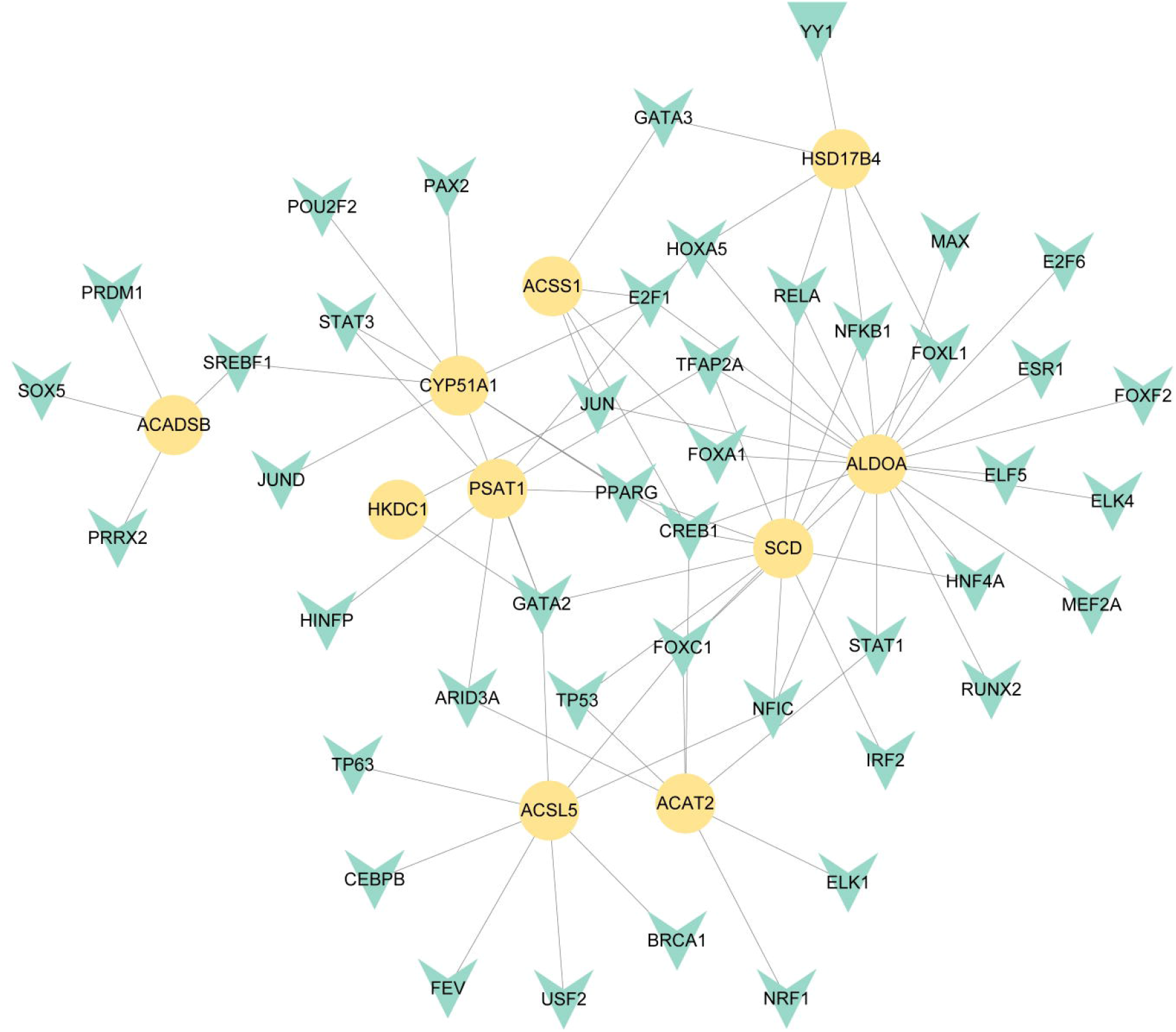
The cohesive regulatory interaction network of hub-gene-TFs obtained from the Network Analyst and described by Cytosacpe. Herein, the green nodes are TFs, and the yellow nodes are hub genes.

**Figure 7.**
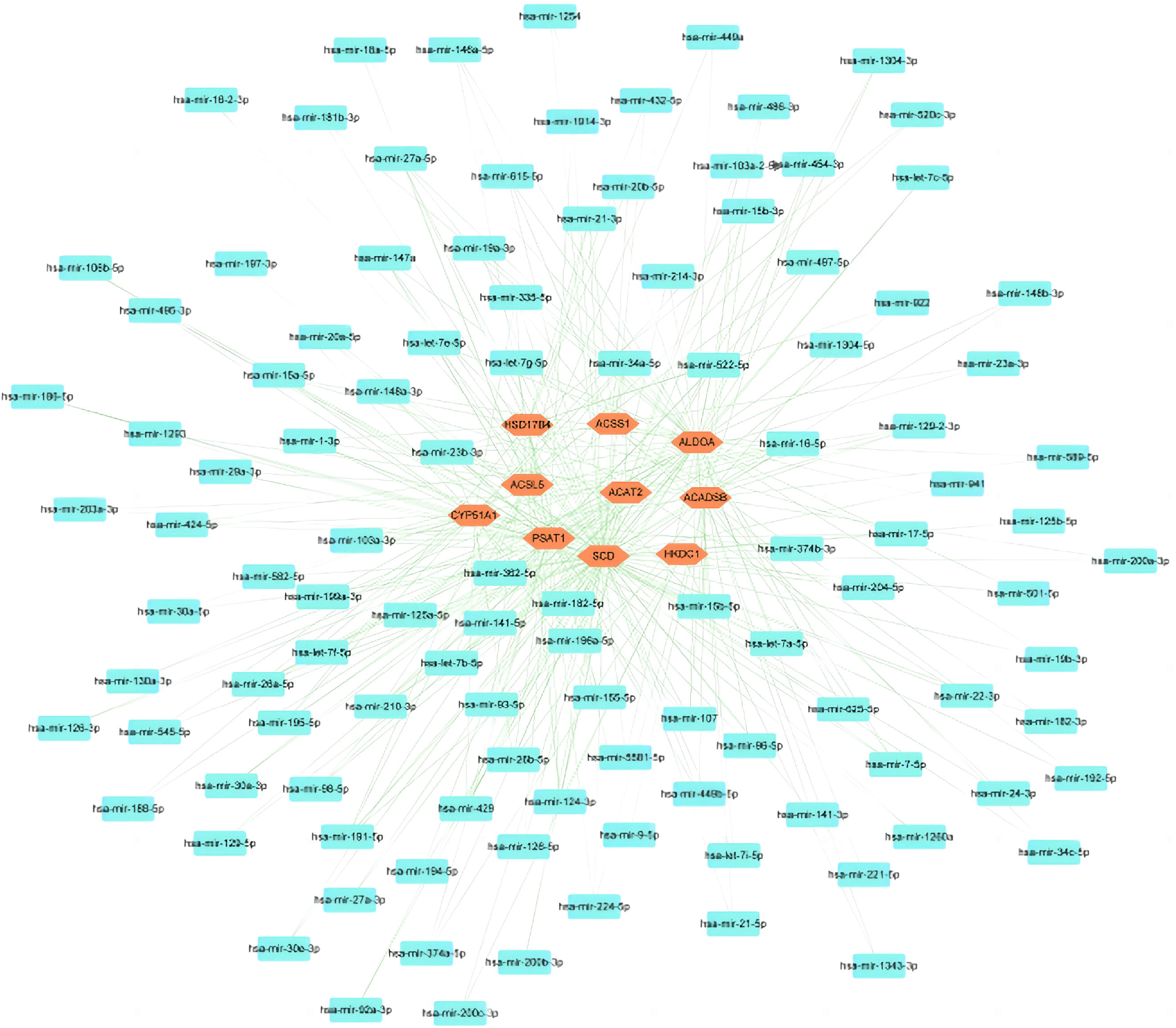
The interconnected regulatory interaction network of hub-gene-miRNAs. Herein, the blue nodes indicate miRNAs and the red nodes are hub genes.

### 3.5 Gene-Disease Association Analysis

If different diseases have one or more similar genes, then we consider these diseases to be related to each other(29). A total of 263 diseases were found to be associated with common genes and screened for significant diseases associated with at least two common genes (Figure 8). In our network, many diseases related to liver and cancer have been found, such as cholestasis, elevated hepatic transaminases, fatty liver, liver cirrhosis, liver dysfunction, mammary neoplasms, neoplasm invasiveness, neoplasm metastasis, non-small cell lung carcinoma, and prostatic neoplasms. Besides, the gene-disease association analysis also found some psychiatric disorders, including epilepsy, hyperreflexia, schizophrenia, and cognitive delay. These results portend the common association of COVID-19 and ICC with these diseases.

**Figure 8.**
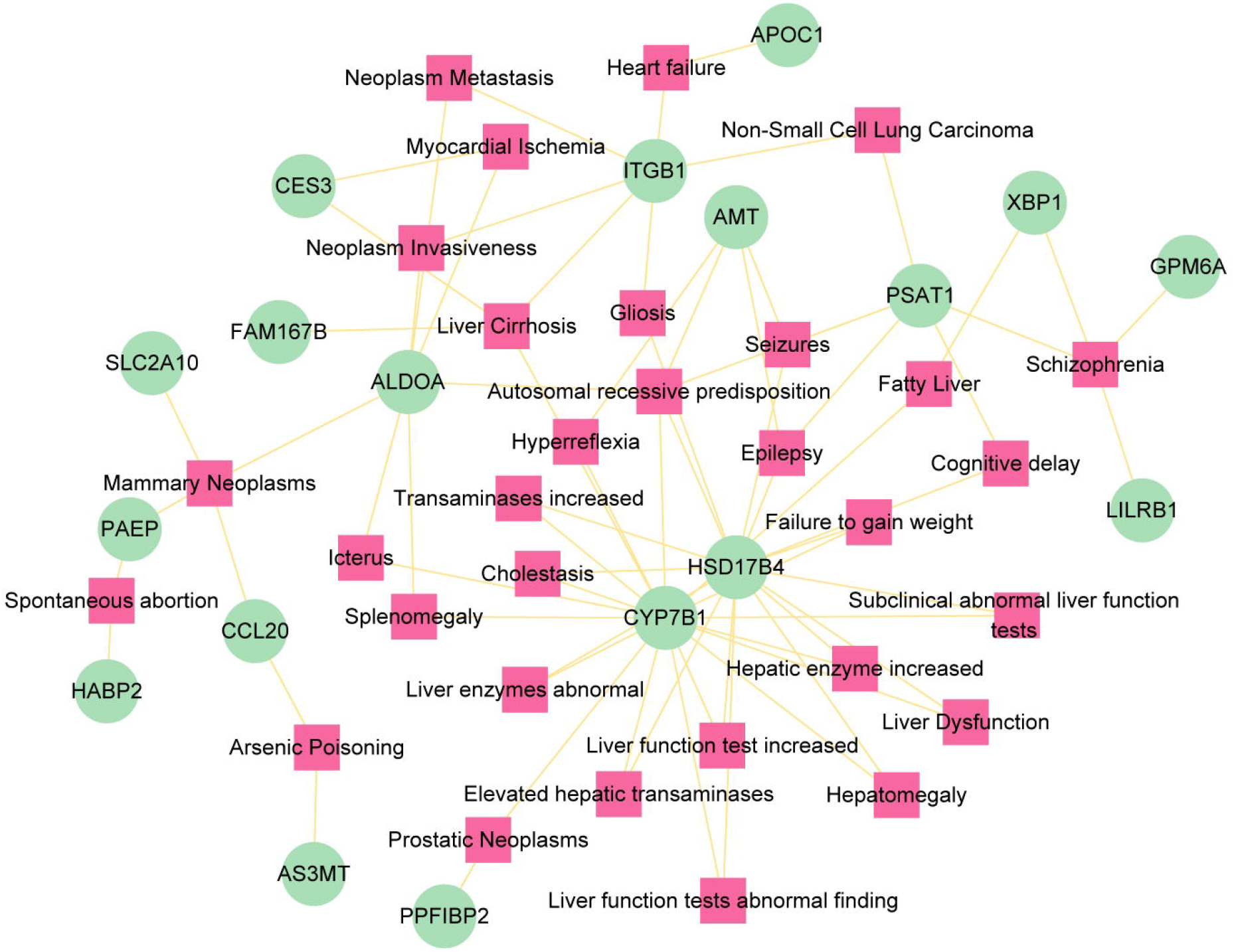
The gene-disease association network represents diseases associated with common genes. The diseases are depicted by the square node and gene symbols are defined by the circle node.

## 4 Discussion

The 2019 SARS-CoV-2 global pandemic has riveted the world’s attention. With more and more variants of the virus, the transmission rate and morbidity rate of COVID-19 gradually increased. Although COVID-19 primarily affects the respiratory system, liver dysfunction is also common in COVID-19 patients, such as elevated liver transaminases and elevations of cholestatic liver enzymes(30). The association between the most common liver cancer, HCC, and COVID-19 has been reported in a previous article(9). ICC, which is the second most common liver cancer and has many similarities with HCC, may also be linked to COVID-19. To verify this idea, bioinformatics analyses were performed on COVID-19 and ICC sequencing data. This study identified 70 shared DEGs and conducted subsequent analyses using these genes, including GO analysis, pathway analysis, PPI network, transcriptional level regulatory networks, and gene-disease network. According to these analyses, GO terms and pathways related to immunity and metabolism were identified. Further, we found hub genes, TFs and miRNAs, which are associated with immunity, metabolism and cancer. Finally, diseases related to hub genes were predicted in this work. This study could establish a link between COVID-19 and ICC and suggest possible treatment options for ICC patients infected with COVID-19.

In our results, lipid biosynthetic process (GO:0008610), glyoxylate and dicarboxylate metabolism pathway, and fatty acid degradation pathway are found in GO terms and pathways. Research uncovered dysregulation of multiple apolipoproteins including APOA1, APOA2, APOH, APOL1, APOD, and APOM in COVID-19 patients. It also detected dysregulated metabolites involved in lipid metabolism in serum(31). Another study also found that SARS-CoV-2 infection elevated the expression of the RE1-silencing transcription factor (REST), which regulated the transcriptional expression of secreted metabolic factors such as myeloperoxidase, apelin, and myostatin, causing disruptions in glucose and lipid metabolism(32). Moreover, recent studies have found that altered lipid metabolism is a new hallmark of cancer(33), as well as ICC. A study found that KDM5C, a histone H3K4-specific demethylase, can repress FASN-mediated lipid metabolism to exert tumor suppressor activity in ICC(34). Consistent with the results of GO and pathway analysis, there are also many genes related to metabolism in hub genes. Stearoyl-CoA desaturase (SCD) was reported to plays a key role in lipid biosynthesis pathways involved in tumorigenesis, and so pharmacological inhibitors have been developed such as MF-438, CAY10566 and A939572(35), but has few research in ICC. In addition, both *ACSL5* and *HSD17B4* were found to be associated with fatty acid synthesis, which may indicate the impact of COVID-19 and ICC on lipid metabolism. These results suggested that COVID-19 and ICC may jointly affect lipid metabolic function of human body. However, whether lipid metabolism can be a therapeutic target for these two diseases needs further study.

Regulation of dendritic cell apoptotic process (GO:2000668), negative regulation of leukocyte apoptotic process (GO:2000107), positive regulation of GTPase activity (GO:0043547), and positive regulation of lymphocyte migration (GO:2000403) are related to immunity, which suggested that both ICC and COVID-19 have a huge impact on the immune system. SARS-CoV-2 has been demonstrated to alter normal immune responses, resulting in a weakened immune system and uncontrolled inflammatory reactions in COVID-19 severe and critical patients(36), which is the major cause of acute respiratory distress syndrome (ARDS). Plasma levels of IL-2, IL-7, IL-10, granulocyte colony-stimulating factor (G-CSF), IP-10, MCP1, macrophage inflammatory protein 1α (MIP1α), and tumor necrosis factor (TNF) have been observed in patients with severe COVID-19 were higher than in healthy adults(37). On the other hand, cancer is usually associated with immune escape by suppressing the immune system. A study found that tumor-derived exosomal miR-183-5p up-regulates PD-L1-expressing macrophages to foster immune suppression and disease progression in ICC through the miR-183-5p/PTEN/AKT/PD-L1 pathway(38). Additionally, in this study, complement and coagulation cascades related pathways are found in top 10 pathway in each database. It has been shown that SARS-CoV-2 may activate the complement system’s classical and lectin pathways(39), and lectin pathways components were found deposited in lung tissue of COVID-19 patients(40), which is consistent with the results of our pathway analysis. Meanwhile, the complement system may be involved in liver dysfunction in viral-induced acute liver failure cases(41). The aforementioned hub protein SCD also plays a role in immune function. A recent study found that suppression of SCD reduces humoral immune response to immunization and weakens immune defense against respiratory influenza infection(42). But SCD1 expressed in cancer cells and immune cells causes immune resistance conditions, and its inhibition augments antitumor T cells and therapeutic effects of anti-PD-1 antibody(43). Not only that, hub gene *PSAT1* can also enhance immunosuppressive through PERK-ATF4-PSAT1 axis in tumor(44, 45). CREB1, a TF with the highest correlation score in our TF-hub gene interaction analysis, was reported to promote T cell cytotoxicity(46). In conclusion, both COVID-19 and ICC can elicit immune system responses. COVID-19 usually causes elevated inflammatory immune response, while ICC causes immune suppression. But the combined effect of these two immune responses on human body is unknown.

There was a report that a patient diagnosed with advanced Hodgkin’s lymphoma, who was not being treated for lymphoma, contracted COVID-19 and four months after ending treatment for COVID-19, was re-examined by PET-CT and found that most of his tumors had disappeared, with levels of biomarkers associated with the tumor dropping by more than 90%(47). Interestingly, the associations between COVID-19 and cancer were also identified in our study. The hub gene in our analysis *ACSL5, ALDOA*, and *HKDC1* are directly associated with liver cancer(48–50). Besides, gene-disease network analysis found some cancer related diseases, including mammary neoplasms, non-small cell lung carcinoma, and prostatic neoplasms. At the same time, neoplasm invasiveness and neoplasm metastasis are also showing in the result, which suggested that the ICC patient with COVID-19 may have a risk of developing other types of tumors and metastases. Similarly, in TF-gene network, SREBF1 was found to enhance the viability and motility in cancer(51). The above evidences suggest that COVID-19 may have an effect on tumor migration and metastasis in ICC patients, but the detailed effect and mechanism require further investigation.

In conclusion, our analysis suggested that COVID-19 and ICC can affect the metabolism and immune system of human body, and may cause the development of new tumors and the metastasis of existing tumors. This study can provide a new perspective for COVID-19 treatment.

## 5 Author Contributions

Tengda Huang, and Kefei Yuan conceived the project and design the protocol; Xinyi Zhou, Tengda Huang, Hongyuan Pan, Jiang Lan, Tian Wu, Ao Du, and Yujia Song performed the data analysis; Xinyi Zhou, Tengda Huang, Yue Lv, and Kefei Yuan wrote the manuscript. All authors contributed to the article and approved the submitted version.

## 6 Funding

This work was supported by grants from the Science and Technology Major Program of Sichuan Province (2022ZDZX0019), the National multidisciplinary collaborative diagnosis and treatment capacity building project for major diseases (TJZ202104), the Natural Science Foundation of China (82272685, 82202260, 82173124, 82173248, 82103533, 82002572, 82002967, 81972747 and 81872004), the fellowship of China National Postdoctoral Program for Innative Talents (BX20200225, BX20200227), the Project funded by China Postdoctoral Science Foundation (2022TQ0221, 2021M692278, 2020M673231), the Science and Technology Support Program of Sichuan Province (2021YJ0436), the Postdoctoral Science Foundation of Sichuan University (2021SCU12007), the Postdoctoral Science Foundation of West China Hospital (2020HXBH075, 2020HXBH007), and the Sichuan University postdoctoral interdisciplinary Innovation Fund (10822041A2103).

## 7 Acknowledgements

We thank all patients who participated in this study and donated samples and the database GEO for providing their platform.

## 9 Supplementary Material

Supplementary Table 1: Differentially expressed genes in GSE119336

Supplementary Table 2: Differentially expressed genes in GSE152418

Supplementary Table 3: Shared DEGs between GSE119336 and GSE152418

Supplementary Table 4: GO terms of DEGs

Supplementary Table 5: KEGG 2021 human pathway, Wikipathway 2021, Reactome 2022 pathway, and the Bioplanet 2019 pathway of DEGs

Supplementary Table 6: PPI network of common DEGs

Supplementary Table 7: TF-Gene topology table

Supplementary Table 8: miRNA-Gene topology table

## Notes

### Competing Interest Statement

The authors have declared no competing interest.

